# Sonic hedgehog is basolaterally sorted from the TGN and transcytosed to the apical domain involving Dispatched-1 at Rab11-ARE

**DOI:** 10.1101/2021.11.10.468081

**Authors:** Lisette Sandoval, Mariana Labarca, Claudio Retamal, Paula Sanchez, Juan Larraín, Alfonso González

**Affiliations:** Centro de Biología Celular y Biomedicina (CEBICEM), Facultad de Medicina y Ciencia, Universidad San Sebastián, Santiago 7510157, Chile; Centro de Envejecimiento y Regeneración (CARE), Facultad de Ciencias Biológicas, Pontificia Universidad Católica de Chile, Santiago 8330025, Chile; Fundación Ciencia y Vida, Santiago 7780272, Chile

**Keywords:** Hedgehog, polarity, trafficking, dispatched, Rab11, lipidation

## Abstract

Hedgehog (Hh) secretion from apical and/or basolateral domains occurs in different epithelial cells impacting development and tissue homeostasis. Palmitoylation and cholestyrolation attach Hh proteins to membranes and Dispatched-1 (Disp-1) promotes their release. How these lipidated proteins are handled by the complex secretory and endocytic pathways of polarized epithelial cells remains unknown. We show that MDCK cells address newly synthesized sonic hedgehog (Shh) from the TGN to the basolateral cell surface and then to the apical domain through a transcytosis pathway that includes Rab11-apical recycling endosomes (Rab11-ARE). Both palmitoylation and cholestyrolation contribute to this sorting behavior, otherwise Shh lacking these lipid modifications is unpolarized. Disp-1 mediates first basolateral secretion from the TGN and then transcytosis from the Rab11-ARE. At steady state, Shh predominates apically and can be basolaterally transcytosed. This complex Shh trafficking provides several steps for regulation and variation in different epithelia, subordinating the apical to the basolateral secretion.

## INTRODUCTION

Hedgehog proteins (Hhs) are signaling determinants of embryonic patterning, differentiation and organogenesis and after birth contribute to tissue homeostasis and repair (Beachy et al., 2004; Briscoe and Therond, 2013; Guerrero and Kornberg, 2014; Parchure et al., 2018; Petrov et al., 2017). Alterations of Hhs functions are involved in developmental defects, tissue fibrosis and cancer (Briscoe and Therond, 2013; Edeling et al., 2016; Machado and Diehl, 2017). In most of these processes the function of Hhs depends on polarized secretion from epithelial cells that possess apical and basolateral domains facing opposite compartments of the organism (Briscoe and Therond, 2013; Guerrero and Kornberg, 2014; Mao et al., 2018; Matusek et al., 2020; Parchure et al., 2018; Peng et al., 2015; Petrov et al., 2017; Walton and Gumucio, 2021). Apical secretion sends signals to neighboring epithelial cells while basolateral secretion targets neighboring epithelial cells and underlying stromal cells (Chamberlain et al., 2008; Lehmann et al., 2020; Mao et al., 2018; Walton and Gumucio, 2021). Elucidating the Hhs intracellular trafficking pathways in epithelial cells is then required to fully understand the signaling function of these important morphogens in development, tissue homeostasis and pathology.

Hhs are synthesized in the endoplasmic reticulum (ER) as a 45kD precursor that is autocatalytically cleaved and processed, generating a ~20kDa N-terminal peptide modified by the covalent addition of cholesterol to its C-terminus and palmitate to its N-terminal end (Chen et al., 2011; Pepinsky et al., 1998; Porter et al., 1996). These lipid moieties attach the mature and functionally competent Hhs to the membrane (Petrov et al., 2017). Multiple forms of Hhs secretion have been described, including oligomers, lipoproteins, exosomes and a main releasing mechanism from the plasma membrane involving the transmembrane and cholesterol binding protein Dispatched (Disp) (Burke et al., 1999; Caspary et al., 2002; Creanga et al., 2012; Parchure et al., 2018; Stewart et al., 2018; Tukachinsky et al., 2012). Disp functions as Hhs/Na+ antiporter driven by the Na+ gradient (Petrov et al., 2020; Wang et al., 2021) and becomes functionally competent after its cleavage at the cell surface by the proprotein convertase Furin, which in epithelial cells occurs at the basolateral plasma membrane (Hall et al., 2019; Stewart et al., 2018).

In different epithelial cells Hhs can be found localized to the apical, basolateral or both plasma membrane domains and their polarized secretion can be inferred from target cell responses. Sonic hedgehog (Shh), the most studied vertebrate orthologs of Hhs (Parchure et al., 2018; Petrov et al., 2017), generates paracrine responses of stromal mesenchymal cells underneath the epithelial cells of adult intestine (Buller et al., 2012; Shyer et al., 2015), lung (Mao et al., 2018; Peng et al., 2015), and kidney (Ding et al., 2012; Edeling et al., 2016; Zhou et al., 2014), indicating basolateral secretion. Instead, gastric parietal cells distribute and secrete Shh mostly apically, showing also detectable levels at the basolateral side (Zavros et al., 2008). Apically secreted Shh is also found in the neural tube of chick (Danesin et al., 2006) and mouse (Chamberlain et al., 2008) embryos. In the mature airway epithelia, Shh is secreted from both apical and basolateral domains, eliciting different signaling pathways and functional consequences at each compartment (Mao et al., 2018).

The routes followed by Hhs along the complex vesicular trafficking pathways of epithelial cells have been mainly explored in *Drosophila* (Guerrero and Kornberg, 2014; Therond, 2012). Extensive studies interpreting subcellular localizations in producing cells and responses of target cells describe both apical and basolateral pools of Hh and propose a primary biosynthetic pathway to the apical domain, followed by endocytosis and either apical recycling (D’Angelo et al., 2015) or transcytosis of Hh to the basolateral domain (Callejo et al., 2011). These alternative pathways remain highly debated and would determine different secretion and signaling characteristics (Gore et al., 2021; Matusek et al., 2020; Simon et al., 2016). How extensive these *Drosophila* models are to mammalian epithelial cells is unknown.

The biosynthetic routes of different epithelial cells display variations reflecting plasticity in the polarity program (Rodriguez-Boulan and Macara, 2014). Exocytic pathways emerging from the TGN can predominate towards the basolateral domain, from where apical proteins then undergo transcytosis, as in intestinal cells and hepatocytes (Bartles et al., 1987; Hauri and Matter, 1991; Rindler and Traber, 1988; Tuma and Hubbard, 2003). In other epithelial cells, epitomized by kidney epithelial cells, apical and basolateral proteins are first sorted at the TGN into distinct polarized secretion pathways of equivalent transport capacity and endocytic proteins can then recycle to the same or the opposite cell surface (Gonzalez and Rodriguez-Boulan, 2009; Mellman and Nelson, 2008; Rodriguez-Boulan and Macara, 2014; Tuma and Hubbard, 2003). Protein trafficking along polarized vesicular pathways is controlled by sorting signals embedded in the cargo and decoded by a complex machinery coupled to the generation of vesicular/tubular vehicles (Rodriguez-Boulan and Macara, 2014). Transmembrane proteins can have apical sorting signals in extracellular, membrane-spanning or cytosolic domains, or basolateral signals located in their cytosolic domains (Bonifacino, 2014; Gonzalez and Rodriguez-Boulan, 2009; Mellman and Nelson, 2008; Rodriguez-Boulan and Macara, 2014). Luminal proteins anchored to membranes through lipid moieties, such as Hh proteins and glycosylphosphatidylinositol-anchored proteins (GPI-APs), associate with lipid rafts (Long et al., 2015; Parchure et al., 2018; Petrov et al., 2017), originally proposed as platforms for apical sorting (Simons and Ikonen, 1997). Although most GPI-APs are addressed to the apical cell surface, there are also examples of basolateral sorting of GPI-APs associated with lipid-rafts (Imjeti et al., 2011; Lebreton et al., 2021; Lebreton et al., 2019; Paladino et al., 2015; Sezgin et al., 2017). Previous studies describe Hhs subcellular distributions and secretion at steady-state conditions that do not define the trafficking pathways followed by newly synthesized Hhs proteins in epithelial cells.

Madin-Darby canine kidney (MDCK) cells forming polarized monolayers in culture constitute the most characterized model system for studies of protein trafficking in epithelial cells (Rodriguez-Boulan and Macara, 2014). In MDCK cells, proteins lacking polarity sorting signals are distributed unpolarized, reflecting TGN-derived apical and basolateral routes of equivalent capacity (Gonzalez et al., 1987; Gottlieb et al., 1986), whereas domain-selective secretion implies specific sorting events occurring at the TGN (Gonzalez et al., 1993; Scheiffele et al., 1995). A unique study in transfected MDCK cells describes Shh steady-state apical and basolateral distribution and enhanced basolateral secretion by coexpressed Disp-1 (Etheridge et al., 2010). This study leaves uncertainty as to whether Shh basolateral secretion occurred directly from the TGN or indirectly through transcytosis from the apical domain, as suggested in Drosophila (Callejo et al., 2011).

A combination of experimental approaches previously used for GPI-APs (Paladino et al., 2006; Zurzolo et al., 1993), transmembrane (Cancino et al., 2007; Gan et al., 2002; Kreitzer et al., 2000) and secretion (Gonzalez et al., 1987) proteins can be applied to define the biosynthetic and endocytic trafficking routes of newly synthesized Shh. Here we evaluate Shh trafficking in MDCK cells using pulse-chase domain-specific targeting assays (Cancino et al., 2007; Paladino et al., 2006), plasmid microinjection expression combined with synchronized post-TGN trafficking imaging of Shh, co-expression with Disp1 in live cells (Cancino et al., 2007; Kreitzer et al., 2003), and cell surface tagging with biotin or antibody. The results reveal that newly synthesized Shh is first sorted from the TGN to the basolateral plasma membrane and then undergo transcytosis to the apical cell surface involving the Rab11-apical recycling endosome (Rab11-ARE). Shh polarized sorting requires the lipid modifications and therefore depends on membrane association. At steady state, Shh is mainly localized at the apical cell surface and its secretion is also mainly apical. A small proportion recycles back to the basolateral cell surface. We also show that Disp-1 first promotes basolateral secretion of newly synthesized Shh transported from the TGN and then is required for its transcytosis through the Rab11-ARE to the apical domain. Such a complex trafficking behavior predicts that apical Shh secretion reciprocally depends on previous basolateral secretion and transcytosis processes that are both sensitive to Disp-1 cleavage and expression levels.

## RESULTS

### Predominant apical distribution and secretion of Sonic Hedgehog in polarized epithelial MDCK cells

Previous studies indicate that Shh can be found distributed at and secreted from apical and basolateral domains with variable preponderance in different epithelial cells (Briscoe and Therond, 2013; Chamberlain et al., 2008; Guerrero and Kornberg, 2014; Mao et al., 2018; Petrov et al., 2017; Shyer et al., 2015; Zavros et al., 2008; Zhou et al., 2014). To explore the trafficking behavior of Shh in a widely characterized epithelial model system (Mellman and Nelson, 2008; Rodriguez-Boulan and Macara, 2014), we generated permanently transfected MDCK cells expressing different levels of Shh and first assessed the polarity of Shh distribution and secretion at steady state conditions. Domain-specific cell surface biotinylation of MDCK-Shh cells grown in Transwell chambers revealed Shh either highly polarized at the apical domain or unpolarized, depending on its expression levels (Figure 1A). For instance, MDCK-Shh clone-1 distributed Shh mainly apically whereas the MDCK-Shh clone-2 displaying higher expression levels displayed Shh similarly at apical and basolateral cell surfaces. E-cadherin and Na-K-ATPase, two basolateral proteins, demonstrated that both MDCK clones are polarized (Figure 1A). Treatment of MDCK-Shh clone 1 with Na^+^-butyrate treatment, known to enhance the expression of transfected plasmids (Burgos et al., 2004; Gonzalez et al., 1987), led to a loss of the predominant apical Shh location, while maintaining the polarity of endogenous E-cadherin Na^+^-K^+^-ATPase (Figure 1A). These results indicate a predominant apical sorting of Shh through a process that can be saturated at high expression levels. We then focused on MDCK-Shh clone-1, named afterwards as MDCK-Shh, which mainly distributed Shh apically. A quantitative assessment showed ~80% Shh distribution at the apical cell surface and more than 90% apical secretion of Shh in this clone (Figure 1B). The predominant apical pattern of Shh can be also observed by indirect immunofluorescence (Figure 1C).

**Figure 1.**
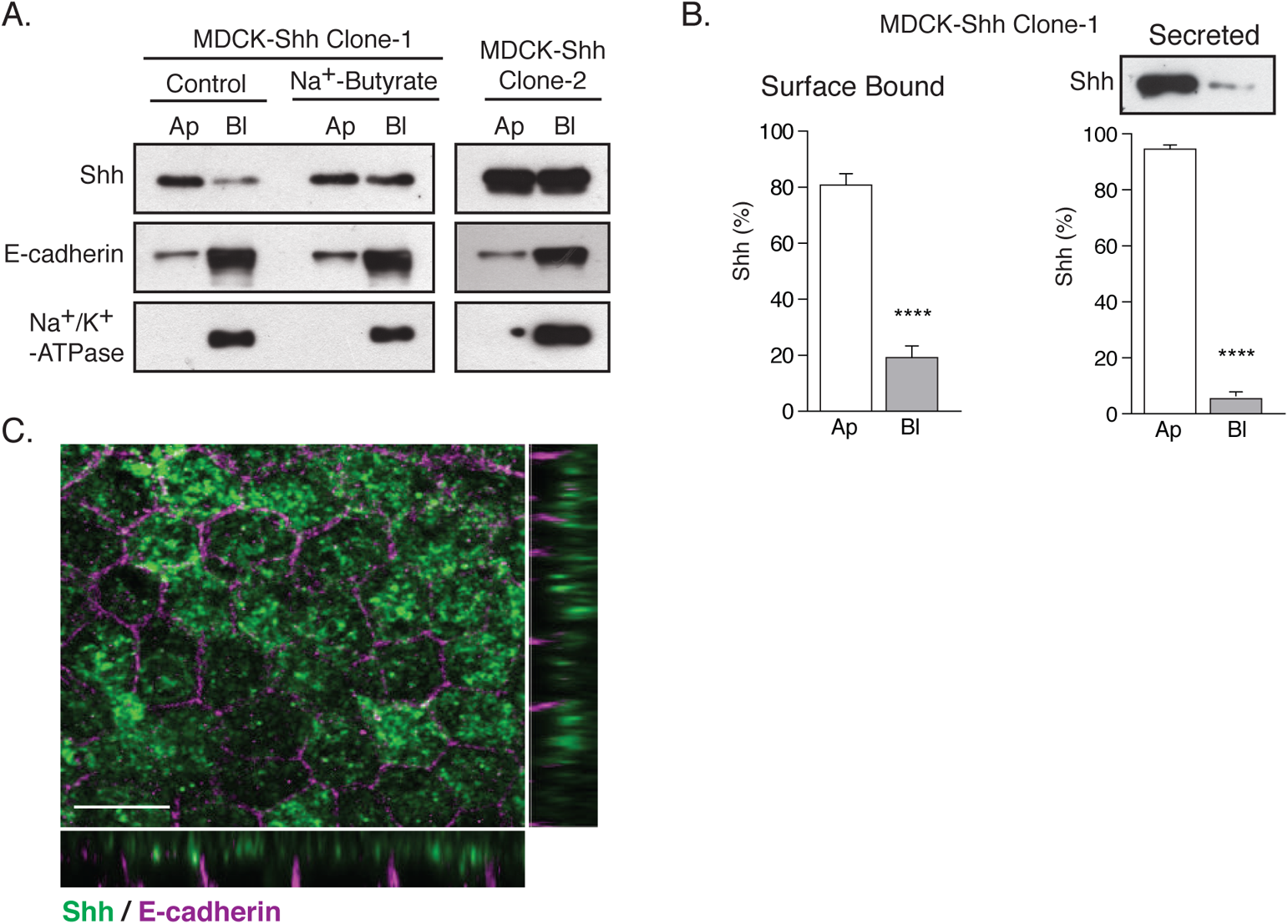
Shh expressed in MDCK cells is apically distributed and secreted. (A) Domain-selective biotinylation of permanently transfected MDCK-Shh grown in Transwell filters. MDCK-Shh Clone 1 shows Shh mainly at the apical cell surface or non-polarized when its expression is increased by 2mM Na+-butyrate treatment for 24h. MDCK-Shh Clone 2 expressing higher Shh levels displays Shh equally distributed among the apical and basolateral cell surface. The basolateral distribution of E-cadherin and Na+/K+-ATPase corroborates the polarized status of the cells in all conditions. (B) Graphs show Shh relative apical versus basolateral cell surface distribution and secretion in MDCK-Shh clone 1. Domain-specific cell surface biotinylation of six independent experiments (n=6) and immunoblot of TCA precipitated proteins from low-serum media conditioned for 6 h of three independent experiments (n=3) reveal aprox. 80% apical cell surface distribution and more than 90% secretion of Shh, respectively. Data expressed as a percentage of total surface or secreted Shh and analysed by an unpaired t-test. In all figures error bars represent mean ± SEM. ****, P≤0.0001. (C) MDCK-Shh clone 1 cells immunofluorescence staining for Shh (green) and E-cadherin (magenta). Images displayed as maximum projection of confocal z-stack and orthogonal views show an apical pattern of Shh. Scale bar, 10 μm.

### Shh is basolaterally sorted and then transcytosed to the apical cell surface

Shh is known to associate with lipid rafts (Long et al., 2015; Parchure et al., 2018; Petrov et al., 2017), similar to GPI-APs, which in general are mainly addressed to the apical domain from the TGN (Hua et al., 2006; Paladino et al., 2006). Therefore, to test whether newly synthesized Shh is directly sorted to the apical cell surface or follows an indirect basolateral-to-apical transcytotic route we pulse-labeled MDCK-Shh cells with [^35^S]-methionine/cysteine and then chased the arrival of newly synthesized [^35^S]-Shh to the apical and basolateral cell surfaces using an established biotinylation targeting assay (Le Bivic et al., 1989; Zurzolo et al., 1992). This approach showed that [^35^S]-Shh first became enriched at the basolateral surface and then progressively increased at the apical cell surface (Figure 2A). E-cadherin used as internal control displayed a basolateral distribution at all chased times. In contrast with Shh, GFP-NO-GPI distributed mostly to the apical cell surface at all chased times (Figure 2B), indicating that our targeting assay is able to detect the TGN-to-apical cell surface pathway previously described for raft-associated GPI-APs in MDCK cells (Imjeti et al., 2011). We also found [^35^S]-Shh in the apical media after 120 minutes of chase, while basolateral secretion was barely detectable (Figure 2C). These results suggest that newly synthesized Shh is primarily sorted to the basolateral domain from the TGN and then is transported by transcytosis to the apical cell surface.

**Figure 2.**
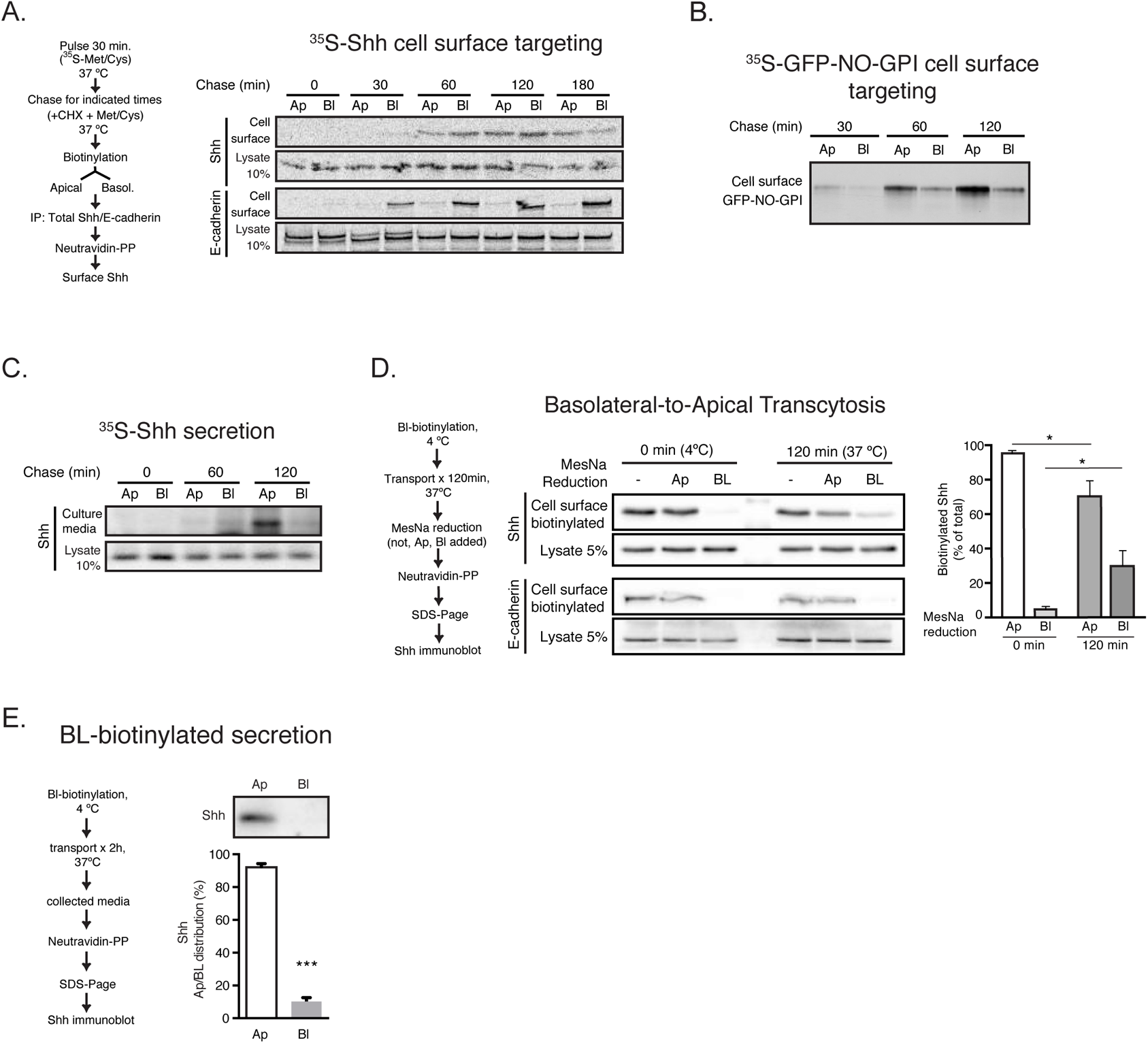
Newly synthesized Shh first arrives at the basolateral cell surface and then reaches the apical side by transcytosis for secretion. Polarized monolayers grown in Transwell filters of MDCK-Shh (A, C) or GFP-NO-GPI (B) cells were pulse-labeled with ^35^S-methionine/cysteine and chased at 37⁰C for the indicated times by subsequent cell-surface-specific biotinylation at 4⁰C. Cell surface targeting of newly synthesized proteins (Shh, E-cadherin or GFP-NO-GPI) and loading control (10% of total protein in lysates) was analyzed by SDS-PAGE and Cyclone radioactivity scanner (see details in Materials and Methods). (C) Immunoprecipitation of ^35^S-Shh shows its secretion to either apical or basolateral chambers. (D) MDCK-Shh were incubated with reducible-Biotin by the basolateral domain at 4⁰C, then allowed to resume trafficking by incubation at 37⁰C for 120 min. Biotin was then reduced by either apical or basolateral sides to detect internalization or transcytosis of the pool of Shh initially biotinylated by the basolateral domain. E-cadherin was used as a control of a basolateral resident protein. Data expressed as a percentage of total surface Shh of three independent experiments (n=3) and analyzed by an unpaired t-test. In graph error bars represent mean ± SEM. *, P≤0.05. (E) MDCK-Shh cells were biotinylated from the basolateral side at 4⁰C and then incubated at 37⁰C for 2h. Apical and basolateral conditioned media were collected, precipitated with Neutravidin and analyzed by immunoblot for secreted Shh. Data expressed as a percentage of total secreted Shh of four independent experiments (n=4) and analyzed by an unpaired t-test. In graph error bars represent mean ± SEM. ***, P≤0.0001.

To directly test basolateral-to-apical transcytosis, we first performed an established assay based on the reducible reagent sulfo-NHS-SS-biotin (Burgos et al., 2004; Zurzolo et al., 1992). After 120 min of transport from the biotinylated basolateral cell surface, approximately 25% of Shh became protected from reduction by MesNa added to the basolateral chamber (Figure 2D). MesNa added to the apical side also decreased the biotinylation levels of basolaterally biotinylated Shh, but not E-cadherin (Figure 2D). Furthermore, we also found in the apical media Shh previously biotinylated from the basolateral cell surface (Figure 2E). These results indicate that Shh already present at the basolateral cell surface reaches the apical cell surface and is apically secreted by transcytosis.

We further corroborate these findings and analyze the route of Shh to the apical domain we tagged the basolateral pool of Shh with anti-Shh antibodies at 4°C and then shifted the cells to 37°C for different times. The basolateral pattern of antibody-tagged Shh (0 min) changed within 60 min to a dot-like pattern typical of the apical recycling endosomes (ARE) and then acquired an apical pattern (Figure 3A). Colocalization of Shh with Rab11 indicated that the subapical dot-like compartment actually corresponds to Rab11-ARE (Figure 3B). Therefore, basolaterally sorted Shh follows the well-known transcytotic route widely characterized for the polyimmunoglobulin (PIgR), which includes the Rab11-ARE (Apodaca et al., 1994; Brown et al., 2000; Wang et al., 2000).

**Figure 3.**
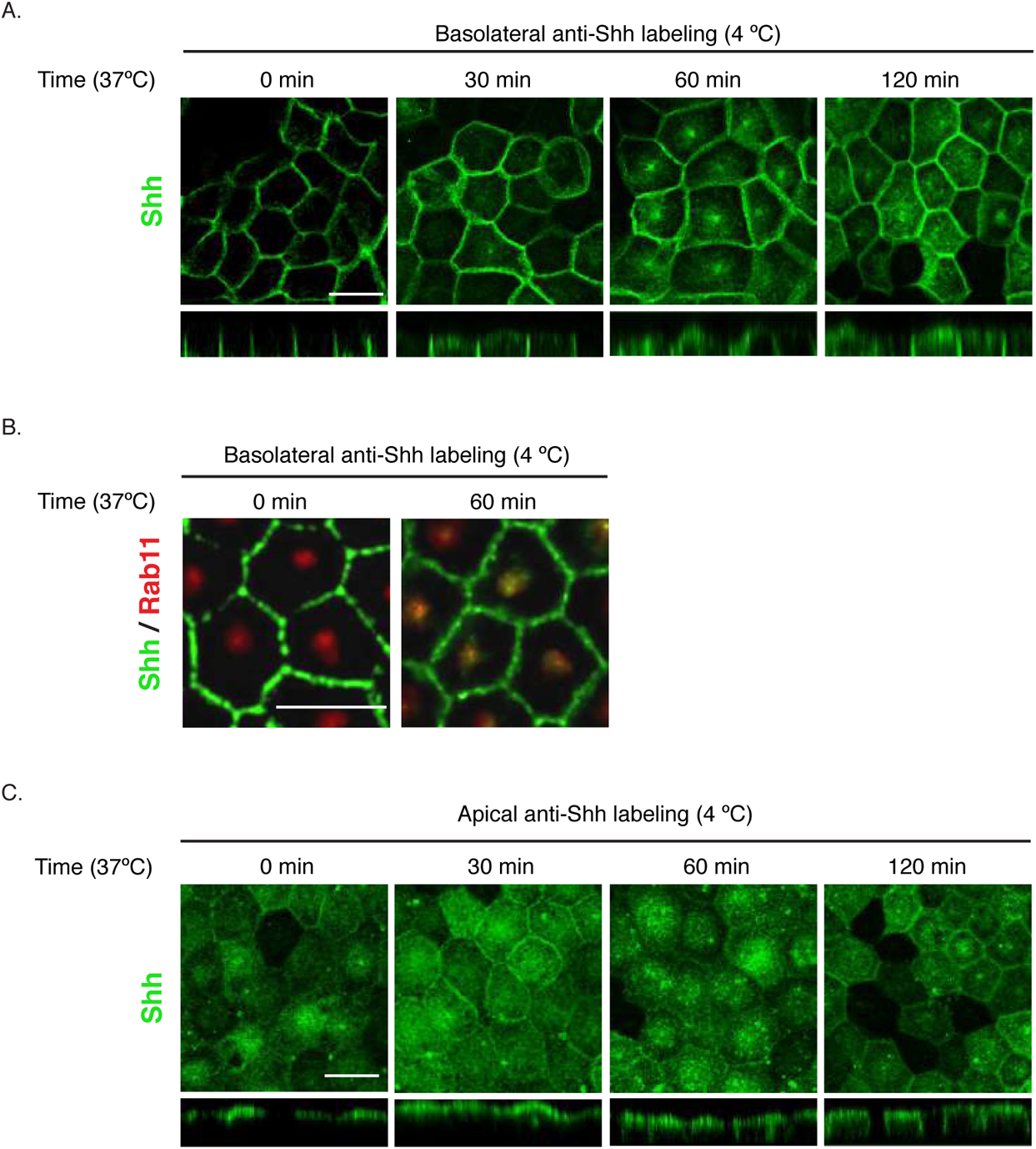
Basolateral Shh moves to the apical side by transcytosis through Rab-11-ARE and apical Shh can undergo reverse transcytosis. MDCK-Shh cells grown in Transwell filters were incubated with anti-Shh antibodies at 4⁰C from the basolateral (A, B) or apical (C) sides, shifted 37⁰C for the indicated times and Shh distribution was followed by immunofluorescence and orthogonal views of confocal images. Shh moved from the basolateral to the apical domain involving a subapical punctate compartment typical of the ARE, as corroborated in by its colocalization with endogenous Rab11 (red) after 60 min of trafficking (B). (C) Apically immunolabeled Shh moving to the basolateral domain is detectable after 60 min. Scale bar, 10 μm.

### Shh can recycle from the apical cell surface back to the basolateral cell surface

Our results so far indicate that newly synthesized Shh follows a transcytotic pathway from the basolateral to the apical membrane. This is congruent with one of the models proposed for Hh protein trafficking in *Drosophila* (Gallet et al., 2003; Gallet et al., 2006). However, the opposite apical-to-basolateral Hh transcytosis has also been proposed in *Drosophila* (Callejo et al., 2011; Guerrero and Kornberg, 2014). Therefore, we used the same type of antibody-tagging experiment and looked for apical-to-basolateral transcytosis. We find that apically tagged Shh can also redistribute to the basolateral cell surface (Figure 3C). Therefore, as suggested in *Drosophila* wing disk epithelia (Callejo et al., 2011; Guerrero and Kornberg, 2014), once arriving to the apical surface Shh can recycle back to the basolateral cell surface by transcytosis.

### Role of lipid modifications in Shh polarized sorting

Hhs proteins are synthesized as a precursor protein that is autocleaved and processed by cholesteroylation at the C-terminus and palmitoylation at the N-terminus (Chen et al., 2011; Pepinsky et al., 1998; Porter et al., 1996) (Figure 4A). These lipid modifications determine Hhs membrane association, nano-scale organization, mode of secretion and range of action (Parchure et al., 2018; Petrov et al., 2017). Therefore, we assessed the polarity of Shh mutants lacking both palmitoylation and cholesterylation (ShhNC24S), only cholesterylation (ShhN) or palmitoylation (ShhNpC24S). ShhNC24S that lacks both lipid modifications and does not associate with membrane (Feng et al., 2004) was secreted without polarity (Figure 4B). Therefore, the peptidic structure of Shh either lacks specific sorting signals for polarity or requires membrane association to expose sorting competent signals, perhaps through nano-scale organization (Vyas et al., 2008; Vyas et al., 2014). We also found the C-terminal truncated mutant ShhN lacking cholesterylation associated to the basolateral cell surface and secreted unpolarized (Figure 4C). Instead, ShhNpC24S lacking the cysteine for palmitoylation and conserving the cholesterol modification preferentially distributed to the basolateral membrane and was mostly secreted apically (Figure 4D). Pulse-chase domain-specific targeting assays showed that newly synthesized ShhNpC24S arrived first at the basolateral cell surface and then at the apical cell surface and medium (Figure 4E), similar to the pathway followed by the fully lapidated protein (See Figure 2A). The low detection levels at the apical cell surface suggest a faster release of this mutant. The transcytosis assay corroborated an apical secretion of basolaterally biotinylated-ShhNpC24S (Figure 4F). These results suggest that ShhNpC24S, similar to wild-type Shh, is initially sorted to the basolateral cell surface and then becomes transcytosed to the apical domain for secretion. All these observations reveal that both palmitoyl and cholesteroyl modifications contribute to Shh sorting to the basolateral cell surface. The cholesterol adduct would be further required for subsequent transcytosis.

**Figure 4.**
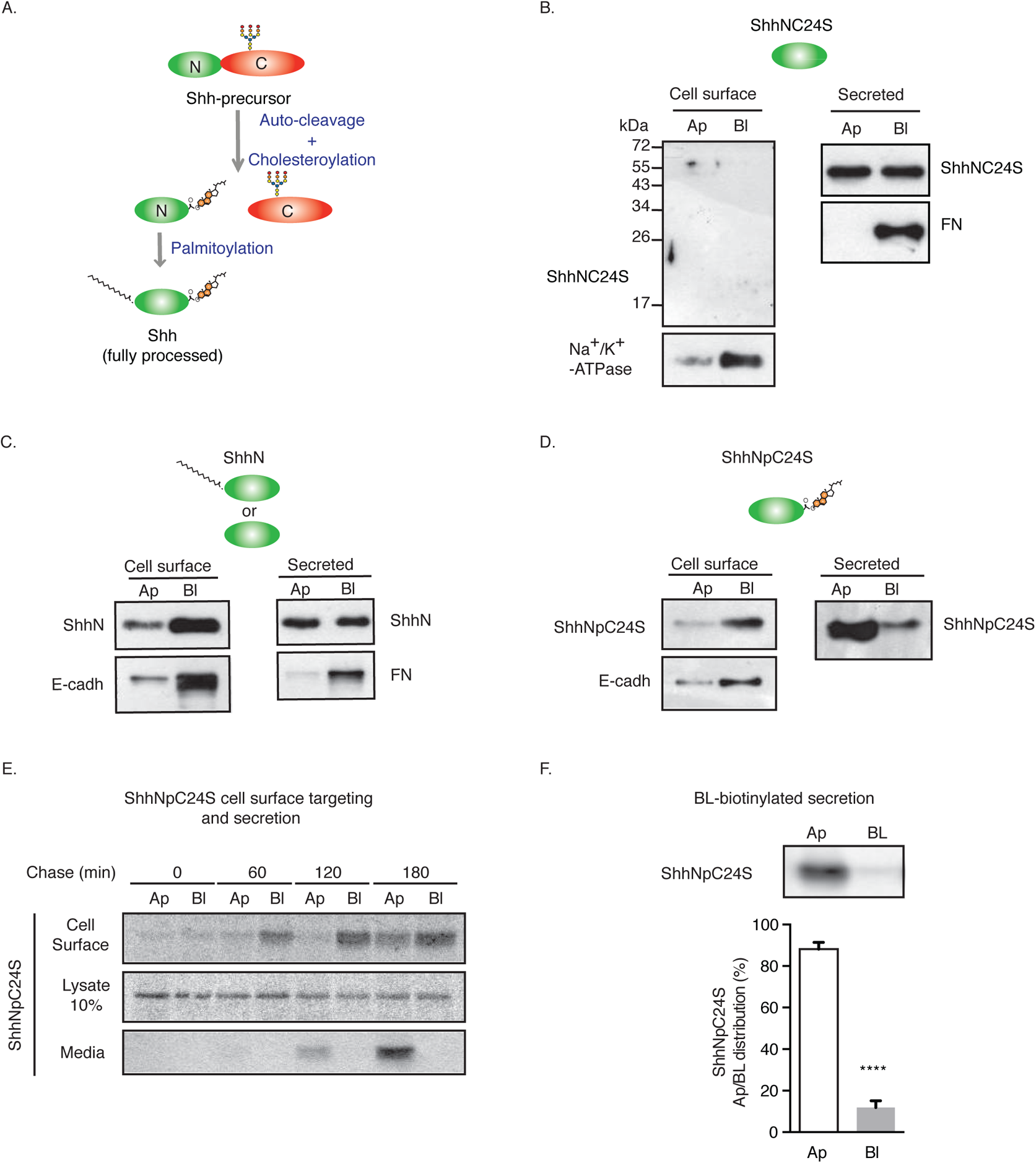
Sorting of Shh lipidation mutants. (A) Schematic illustration of Shh processing and lipidation. (B-F) MDCK cells stably expressing Shh lipidation mutants lacking both lipids (B, ShhNC24S), containing only palmitoyl (C, ShhN) or cholesteryl (D-F, ShhNpC24S) moieties were analyzed by domain-specific cell-surface biotinylation and secretion (presence in the media after 2h accumulation, TCA precipitation and immunoblot). Basolateral E-cadherin and Na+/K+-ATPase transmembrane proteins and secreted Fibronectin (FN) were used to control polarity. (E) MDCK-ShhNpC24S cells were pulsed labeled with ^35^S-methionine/cysteine and chased to detect protein arrival to each cell surface along with its polarized secretion, as in Figure 2A. Samples were analyzed by SDS-PAGE and Cyclone radioactivity scanner. (F) MDCK-ShhNpC24S cells were biotinylated from the basolateral side at 4⁰C and then incubated at 37⁰C for 2h. To assess for ShhNpC24S secretion, apical and basolateral conditioned media were collected, precipitated with Neutravidin and analyzed by immunoblot. Data expressed as a percentage of total secreted Shh of three independent experiments (n=3) and analysed by an unpaired t-test. In graph error bars represent mean ± SEM. ****, P£0.0001.

### Disp-1 distribution and role in newly synthesized Shh secretion

Disp is required for the release of Hh proteins from expressing cells (Burke et al., 1999; Caspary et al., 2002; Petrov et al., 2017; Stewart et al., 2018). In polarized epithelial cells, Disp has been described to promote Hh basolateral secretion (Callejo et al., 2011; Etheridge et al., 2010; Stewart et al., 2018). We used plasmid microinjection (Cancino et al., 2007; Kreitzer et al., 2003) to express Shh alone or together with HA-tagged Disp-1 (Stewart et al., 2018). Microinjection avoids the high levels of overexpression achieved by transient cotransfection experiments, which can alter polarized sorting (Burgos et al., 2004). After 4 h post-microinjection we clearly found Shh distributed apically, contrasting with the co-expressed basolateral marker LDLR-GFP (Cancino et al., 2007; Gravotta et al., 2007; Guo et al., 2013) (Figure 5A). Using these conditions, we co-expressed Shh together with Disp-1 and incubated the intact cells with anti-Shh added to the apical side to detect Shh at the cell surface. Then we permeabilized the cells for Disp-1 immunodetection and found it localized just below the Shh labeled at the apical cell surface (Figure 5B). In the next experiments, we fixed and permeabilized the cells to immunolocalize Shh and Disp-1 at the same time. Confocal images were then merged and shown as either maximal projections or just a basolateral section at the level of the nucleus (Figure 5C). Shh expressed alone can be seen in its typical apical location Shh (Figure 5C, upper panel), while when co-expressed with Disp-1 also became visible at the basolateral border of neighboring non-microinjected cells (Figure 5C, middle panel). This indicates that Disp-1 enhanced the basolateral secretion of Shh, as previously described (Etheridge et al., 2010). Shh and Disp-1 mostly segregated from each other, with Disp-1 occupying a subapical location overlaid by Shh. Both proteins also colocalized in few spots (Figure 5C, middle panel).

**Figure 5.**
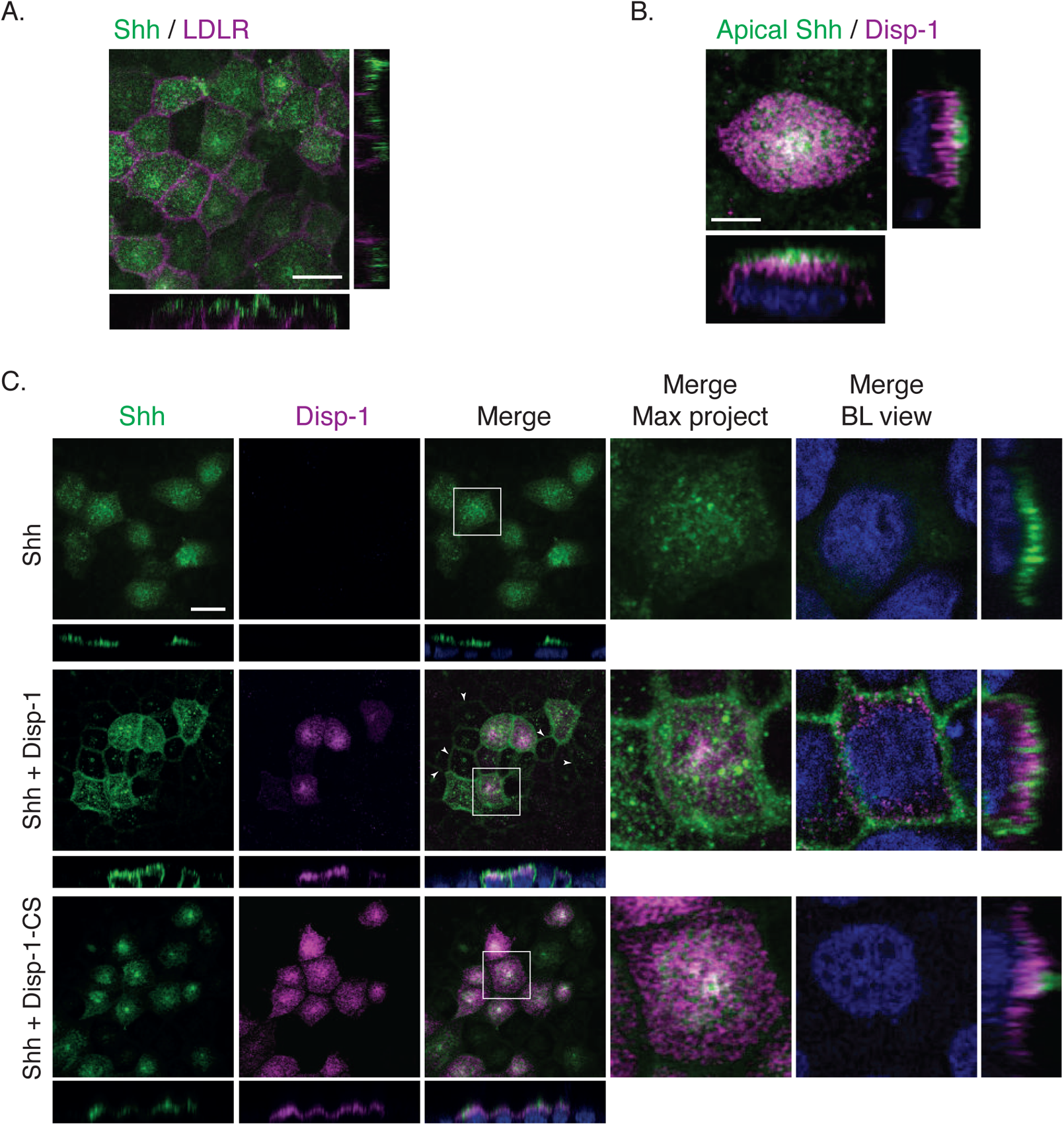
Shh is apically and basolateraly distributed under Dispt-1 expression while it is retained at ARE in cells expressing cleavage deficient Disp-1CS. MDCK cells were plasmid microinjected and protein expression was allowed for 4h. (A) Immunofluorescence of co-microinjected Shh (green) and basolateral marker LDLR-GFP-Y18A (magenta). (B) Co-microinjected cells with Shh and Disp-1 were fixed and immune-stained first by the apical side with anti-Shh (green) antibodies and then permeabilized and stained for Disp-1 with anti-HA (magenta) antibodies. (C) Cells were microinjected with Shh alone (upper panels) or together with Disp-1 (middle panels) or cleavage deficient Disp-1-CS (lower panels). Then cells were analyzed by indirect immunofluorescence for Shh (green) and Disp-1 (HA, magenta). Arrowheads indicate non-microinjected cells with basolateral Shh localization due to secretion from expressing cells. Zoomed image to the right demonstrates Shh apically and basolaterally detected in the presence of Disp-1 (BL view) or retained in the ARE compartment in the presence of Disp-1-CS (maximal projection, also see Fig. 7). In all figures scale bar, 10 μm.

Then, we co-expressed Shh together with the cleavage deficient recombinant mutant Disp-1-R279A/E280A (Disp-1-CS) incompetent for Shh secretion (Stewart et al., 2018). Disp-1-CS displayed an overall similar distribution as Disp-1. However, Disp-1-CS coexpressing cells showed Shh predominantly at the subapical compartment, likely corresponding to the ARE (Figure 5C, bottom panel), and lost the Shh apical location in expressing cells and its basolateral detection in neighboring cells (Figure 5C, bottom panel). These results suggest that Disp-1-CS not only inhibited Shh secretion from the basolateral surface where Shh first arrived, but also arrested Shh at the ARE, while *en route* to the apical cell surface from the basolateral plasma membrane.

Using less amounts of microinjected plasmids to achieve lower levels of expression we detected colocalization of Shh and Disp-1 in the Rab11-ARE compartment, which included tubular structures (Figure 6). High resolution images (3D surface rendering) revealed colocalization of Shh, Disp-1 and Rab11 in the tubular structures of the ARE compartment (Figure 6; middle panel, arrowheads). Even though most of Disp-1 is seen at subapical planes, overexposure of Z confocal planes corresponding to the basolateral volume of the cell clearly showed Disp-1 near the lateral membrane, where few vesicles containing Shh or Rab11 were also detectable (Figure 6, bottom panel).

**Figure 6.**
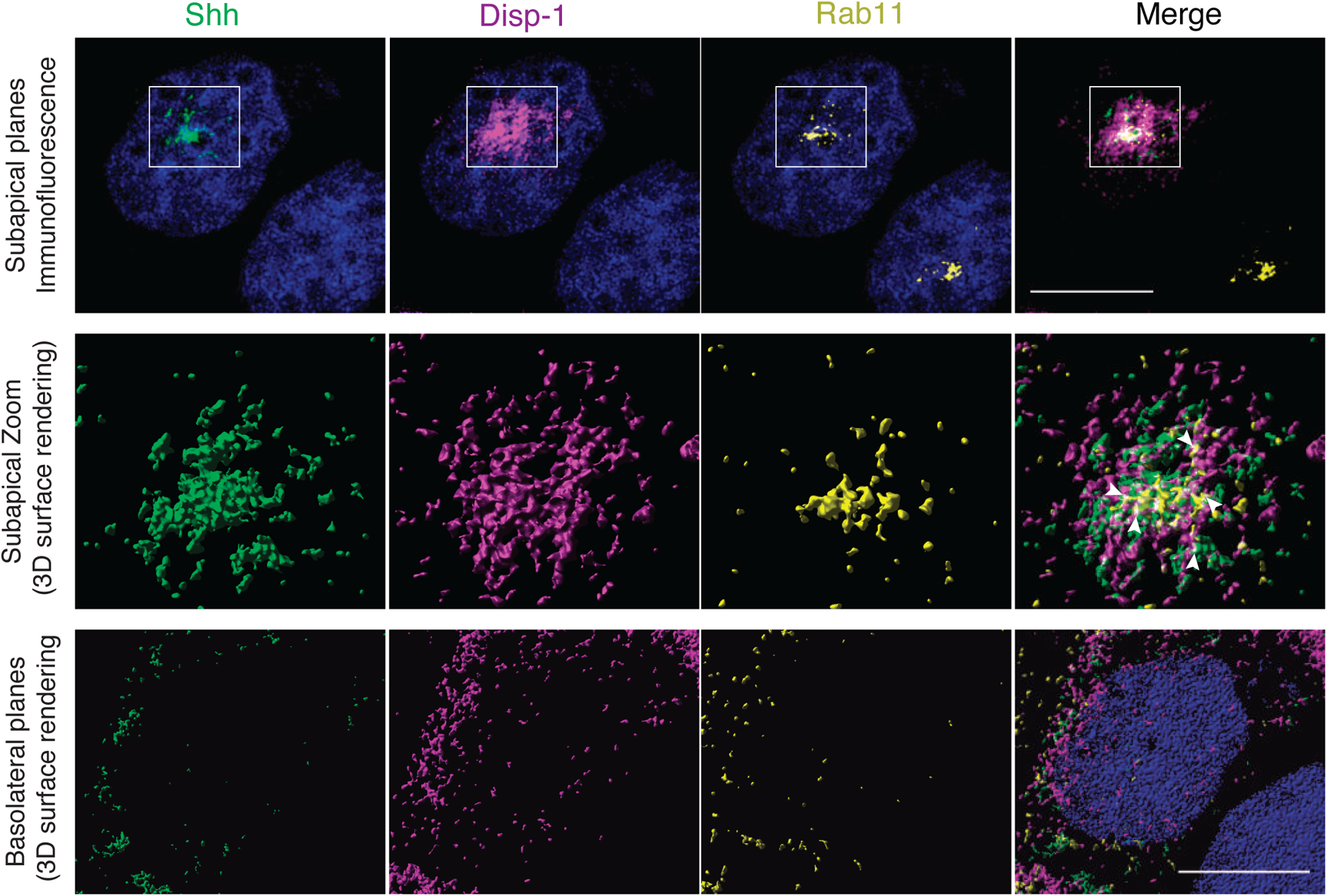
Colocalization of transiently expressed Shh and Disp-1 in Rab11-ARE. Polarized MDCK cells were microinjected with Shh-GFP (green) along with Disp-1 and after 4 hrs expression cells were fixed and immunostained for anti-HA (magenta) and anti-Rab11 (yellow). A microinjected cell is shown in the upper panel (center) and a non-microinjected cell is visible at the down-right corner. Maximal projection of 10 planes of 0.1 microns sections over thshows the 3D surface rendering of the zoomed region, in which colocalization of Shh (green) Disp-1 (magenta) e nucleus shows the subapical Rab11-ARE displaying a tubular pattern. Center panel and Rab11 (yellow), is shown by white arrowheads. The lower panel illustrates a 3D surface rendering of medial and basal planes (40 planes of 0.1 microns sections), in which basal distribution of Disp-1 and few vesicles containing Shh or Rab11 are shown. Scale bar, 10 μm.

To further analyze the role of Disp-1 upon newly synthesized Shh biosynthetic trafficking we performed live-cell imaging from the TGN to cell surface transport of Shh coupled to GFP (Shh-GFP) (Chamberlain et al., 2008). We first corroborated in plasmid-microinjected MDCK cells for Shh-GFP to acquire an apical distribution (Figure 7A) using the route of Rab-11-ARE (Figure 7B). Then, we followed Shh-GFP trafficking from the TGN to the cell surface in a synchronized manner with previously established methods (Cancino et al., 2007; Guo et al., 2013; Kreitzer et al., 2003). After plasmid microinjection, we incubated the cells for 45 minutes at 37°C to allow for detectable expression levels and then incubated the cells at 20°C for 2h to arrest the biosynthetic trafficking at the TGN. At this initial time, Shh-GFP expressed alone or in combination with Disp-1 or Disp-1-CS mutant displayed the perinuclear pattern characteristic of the TGN (Figure 7C, 0 min) (Cancino et al., 2007). Shifting the cells to 37°C allowed the TGN exit in the presence of cycloheximide to avoid further protein synthesis. Shh-GFP first became detectable at the cell border, reflecting its basolateral arrival, and then progressively acquired an ARE location and the typical apical pattern (Figure 7C). A halo of receiving cells surrounding the expressing cells displayed Shh in their border, revealing the Shh basolateral secretion from producing cells (see also Fig 5C middle panel). This can be seen more clearly in the overexposed image corresponding to 120 min post-TGN trafficking (Figure 7C, middle panel). Cells co-expressing Disp-1 showed an enhanced Shh-GFP basolateral secretion, as judged by the higher intensity and longer distance of the fluorescent borders of neighboring non-expressing cells (Figure 7C, middle panel). These results indicate that post-TGN basolateral secretion of Shh is sensitive to Disp-1 expression levels and provide direct evidence of Shh sorting from the TGN to the basolateral cell surface. In striking contrast, cells co-expressing the Disp-1-CS mutant showed a decreased basolateral distribution in neighboring cells (Figure 7C, lower panel), indicating lower basolateral secretion levels. Furthermore, Shh-GFP accumulated in a punctate compartment suggesting an arrest at the ARE (Figure 7C, lower panel). Taken together, all these experiments indicate that Disp1 promotes Shh secretion first from the basolateral cell surface where it arrives from the TGN. The effects of coexpressing Disp-1-CS corroborate that Shh secretion requires Disp-1 processing by cleavage (Stewart et al., 2018) and suggest that Disp-1 is also required for trafficking along the basolateral-to-apical transcytosis route at the level of the ARE, where Shh accumulates if Disp-1 function is disturbed.

**Figure 7.**
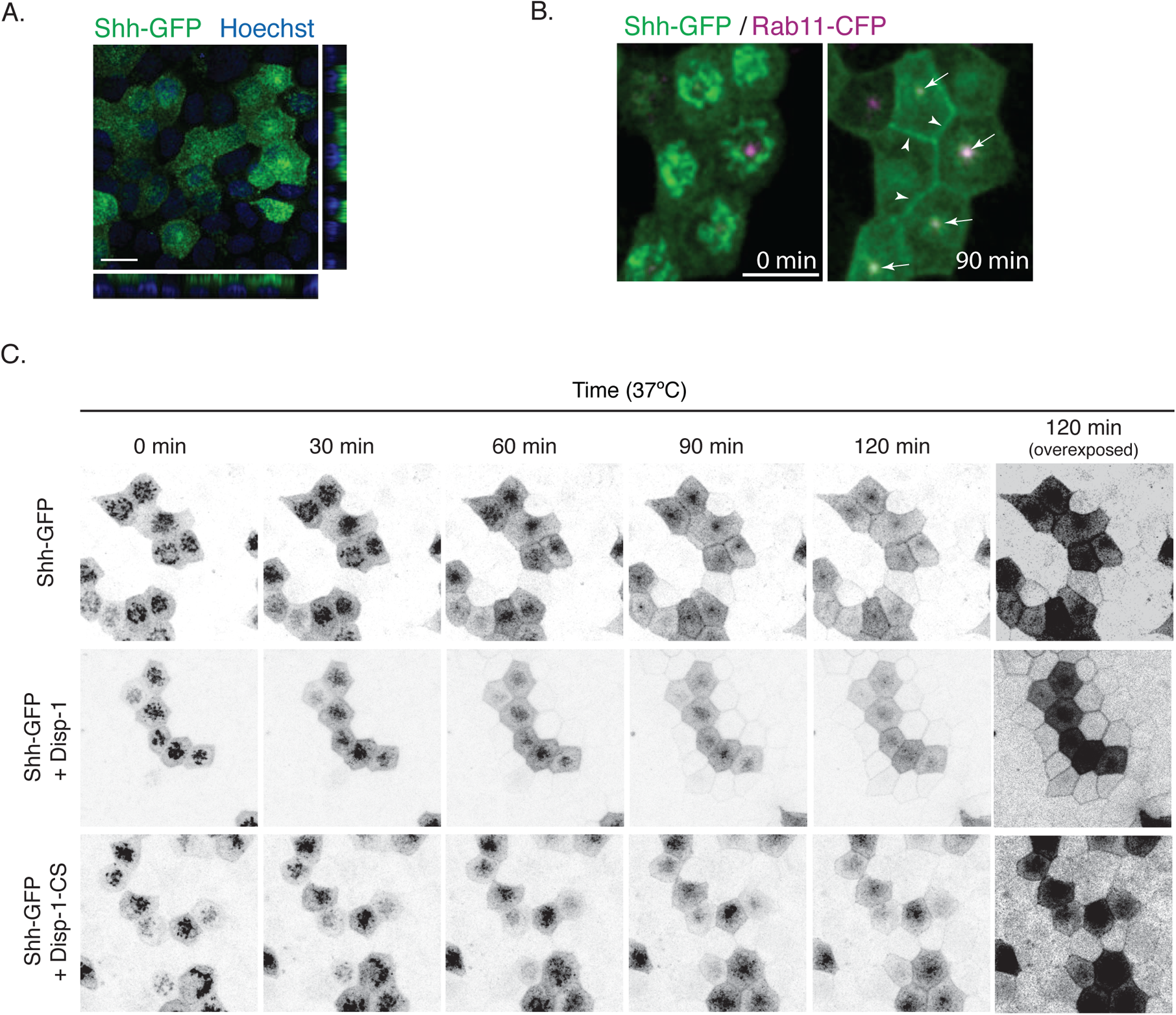
Basolateral secretion of Shh from the TGN is enhanced by Disp-1 and decreased by Disp-1-CS. (A) Shh-GFP distribution after 4 h of expression in plasmid microinjected polarized MDCK cells show a similar polarity as Shh at steady state (See Figure 1). (B) Shh-GFP expressed by microinjection in MDCK cells permanently transfected with Rab11-CFP plasmid. After 45 min of Shh-GFP expression, the cells were incubated for 2 h at 20°C to arrest the protein at the TGN (time = 0) and then shifted to 37°C to reestablish the exit from TGN for 90 min. Arrows point to Rab11-ARE and arrowheads show basolateral Shh-GFP. (C) Live cell imaging analysis of Shh-GFP trafficking from the TGN to the cell surface for the indicated times. Shh-GFP alone (upper panel), co-microinjected with Disp-1 (middle panel) or Disp-1-CS (lower panel). Shh-GFP detected in the border of neighboring non-expressing cells reflects its basolateral secretion, which is enhanced by co-expressed Disp-1 and decreased by Disp-1-CS, more clearly seen in the overexposed image at time 120 min.

## DISCUSSION

In diverse epithelial systems, Hhs are found apically and basolaterally distributed and secreted with variable polarity (Chamberlain et al., 2008; Mao et al., 2018; Therond, 2012). An open question is whether Hh subcellular distribution and secretion reflects default-unsorted or specific-sorting trafficking events. The steady-state conditions of previous studies cannot answer this question and have even led to controversial models (Ayers et al., 2010; Callejo et al., 2011; Gallet, 2011; Guerrero and Kornberg, 2014; Therond, 2012). We addressed this problem using MDCK cells as model system and combined pulse chase and live cell imaging to follow the cell surface arrival of newly synthesized Shh, while domain specific cell surface tagging directly assessed transcytosis from each polarized domain. Our results provide new information regarding the intracellular routes taken by newly synthesized Shh, which can help to interpret and understand previous observations in different epithelial systems.

We find a predominant (~80%) apical cell surface distribution and secretion of Shh in permanently transfected MDCK cells under steady state conditions. However, pulse-chase domain-specific targeting assays reveal that newly synthesized Shh arrives first to the basolateral cell surface and then to the apical plasma membrane. Live cell imaging after releasing a 20C° TGN arrest demonstrates Shh basolateral sorting directly from the TGN. Basolateral tagging with biotinylation or antibody binding directly demonstrate transcytosis of basolateral Shh to the apical domain. The transcytosis pathway of Shh includes the Rab11-ARE compartment, similar to the most characterized transcytosis route of pIgR (Apodaca et al., 1994; Rojas and Apodaca, 2002). A reverse transcytosis of Shh from the apical to the basolateral domain can also occur, as shown by apical tagging with antibody. All these observations suggest a model in which newly synthesized Shh is first addressed to the basolateral domain from the TGN and then becomes transported through the Rab11-ARE transcytotic route to the opposite apical domain, from which most of its secretion takes place while a small proportion returns to the basolateral cell surface. Our results imply specific sorting events operating first at the TGN and then at the basolateral plasma membrane.

MDCK cells possess apical and basolateral exocytic pathways of roughly equivalent transporting capacity and therefore newly synthesized proteins lacking sorting information would follow these pathways by default resulting in their unpolarized distribution or secretion (Gonzalez et al., 1987; Gottlieb et al., 1986; Scheiffele et al., 1995). In contrast, intestinal epithelial cells and hepatocytes have a prominent basolateral exocytic route into which both basolateral and apical proteins can enter and once reaching the cell surface the apical proteins become transcytosed to the opposite domain (Bartles et al., 1987; Hauri and Matter, 1991; Hubbard, 1989; Rindler and Traber, 1988). Proteins lacking information for TGN-mediated sorting that are unpolarized in MDCK cells can be basolaterally addressed in intestinal cells (Rindler and Traber, 1988). Our results in MDCK cells clearly indicate that Shh possess basolateral sorting information decoded first at the TGN and predicts its primary basolateral cell surface arrival in different epithelial cells.

We initially considered that Shh sorting behavior might share similarities with GPI-proteins due to their common association with lipid rafts at the luminal membrane leaflet (Mayor and Riezman, 2004). GPI-proteins are most commonly addressed to the apical cell surface from the TGN in MDCK cells (Imjeti et al., 2011; Lebreton et al., 2019; Lisanti et al., 1988; Paladino et al., 2006). However, our pulse chase experiments revealed basolateral sorting of Shh under the same conditions that corroborated apical biosynthetic trafficking of a previously characterized recombinant GPI-protein (Imjeti et al., 2011). Lipid-raft association is certainly not enough for apical sorting (Lebreton et al., 2019; Paladino et al., 2004). A proposed additional requirement for at least apical GPI-proteins is clustering at the TGN promoted by cholesterol and luminal calcium (Lebreton et al., 2021; Paladino et al., 2004), or glycosylation in some epithelial cells (Imjeti et al., 2011). GPI-proteins lacking this clustering property, such as the prion protein, are instead addressed to the basolateral domain (Lebreton et al., 2021; Sarnataro et al., 2002). Prion also becomes transcytosed from the basolateral to the apical domain (Arkhipenko et al., 2016), similar to what we found here for Shh. The mechanism through which lipid-raft associated proteins are basolaterally sorted, endocytosed and addressed to the opposite domain by transcytosis remains unknown.

Epithelial cells have transcytosis pathways from the apical or basolateral domains to the opposite cell surface (Mostov et al., 1995; Tuma and Hubbard, 2003). Transmembrane proteins can follow these pathways guided by specific sorting signals or associated with default movement of membranes. For instance, the cytosolic tail of pIgR contains sorting signals for basolateral delivery, clathrin-mediated endocytosis and transcytosis (Luton et al., 2009; Mostov et al., 1995), whereas the basolateral recycling transferrin receptor reaches the apical domain when cells do not express the AP1B-adaptor responsible for its basolateral sorting (Perez Bay et al., 2013), suggesting a default post-endocytic trafficking. Both pathways include transit through the Rab11-ARE compartment (Luton et al., 2009; Mostov et al., 1995; Perez Bay et al., 2013). We show that Shh still associated with the basolateral plasma membrane moves towards the apical cell surface using the Rab11-ARE transcytotic route. The mechanism might involve yet unidentified transmembrane escorting proteins linking Shh to the cytosolic sorting machinery or a default pathway from the basolateral cell surface.

Cholesterol modification is a crucial determinant in Hh protein membrane association, mode of secretion and morphogenetic extension range (Callejo et al., 2011; Gallet et al., 2003; Lewis et al., 2001; Porter et al., 1996). We find that Shh mutant lacking palmitoylation and cholesterylation is secreted unpolarized. Therefore, the Shh peptidic structure either lacks specific sorting signals for polarity or requires membrane association to expose sorting competent proteinaceous signals. Our results also show that mutants lacking either palmitoylation (ShhNpC24S) or cholesterylation (ShhN) predominantly distribute to the basolateral cell surface, suggesting that both lipid modifications contribute to the basolateral sorting of Shh. We could not attribute a particular role of palmitoylation in the basolateral-to-apical trafficking because Shh defective in cholesterol modification is inefficiently palmitoylated (Chen et al., 2004; Feng et al., 2004; Kohtz et al., 2001; Sbrogna et al., 2003). On the other hand, our pulse chase and basolateral tagging experiments indicate that the cholesteryl group can drive the entire polarized trafficking pathway of Shh, including the primary basolateral sorting and subsequent transcytosis to the apical domain. It is possible that cholesterol represents a sorting signal decoded at different steps of the Shh itinerary to the apical cell surface, perhaps involving the trafficking role proposed for Disp (Callejo et al., 2011; D’Angelo et al., 2015; Gore et al., 2021; Stewart et al., 2018), as a cholesterol binding protein (Cannac et al., 2020; Chen et al., 2020).

Disp-1 is a multipass transmembrane protein with a luminal/extracellular sterol-sensing domain that binds Shh in a cholesterol-dependent manner and is required for its release in a diffusible form from the membrane (Burke et al., 1999; Cannac et al., 2020; Chen et al., 2020; Creanga et al., 2012; Tukachinsky et al., 2012). In vertebrates, Disp-1 transfers Shh from the cell surface to secreted SCUBE2 requiring the sodium gradient across the plasma membrane (Burke et al., 1999; Cannac et al., 2020; Chen et al., 2020; Petrov et al., 2020). Stewart et al. (Stewart et al., 2018) recently described that Disp-1 acquires functional competence after its cleavage by the proprotein convertase furin at the cell surface (Stewart et al., 2018). Because furin recycles between the TGN and the basolateral cell surface (Simmen et al., 1999), Disp-1 should reach first the basolateral cell surface to become competent in Hhs secretion in epithelial cells (Stewart et al., 2018). In *Drosophila*, exogenously expressed Disp has been reported at the plasma membrane of the apical, basal and lateral regions (D’Angelo et al., 2015), or mainly at the basolateral plasma membrane (Callejo et al., 2011; Stewart et al., 2018). In MDCK cells, transfection experiments showed Disp-1 distributed to the basolateral region seemingly due to sorting information contained in its intracellular domain (Etheridge et al., 2010). We find with RT-PCR that MDCK cells express Disp-1 (not shown) and our microinjection experiments avoiding a saturating overexpression revealed recently synthesized Disp-1 predominantly distributed at the subapical region, just underneath Shh, though also spreading throughout the cell. These observations most likely reflect a dynamic Disp-1 trafficking and variations depending on its expression levels and experimental conditions.

Our experiments of plasmid microinjection combined with live cell imaging show that slightly overexpressed Disp-1 enhances basolateral secretion of newly expressed Shh just after releasing a TGN block. Etheridge et al. (Etheridge et al., 2010) described a decreased basolateral secretion of Shh under conditions of Disp-1 silencing or Disp-1 mutant expression in MDCK cells. Accordingly, we find that coexpression of the cleavage defective Disp-CS mutant (Stewart et al., 2018) decreases the basolateral secretion of Shh. A trade-off between apical and basolateral Hh secretion has been previously proposed in the wing disk epithelia (Ayers et al., 2010; Gore et al., 2021). Our results showing that basolateral secretion of Shh directly from the TGN requires and can be enhanced by increasing the coexpressed level of Disp-1 suggesting that conditions that increase or decrease this process would determine how much Hhs reaches the apical cell surface, where Disp is also expected to mediate Hhs secretion, (D’Angelo et al., 2015).

Disp has been proposed to play a role in Hh subcellular distribution and trafficking through endocytic pathways. This notion derives from studies in Drosophila epithelia models where Hh secretion and signaling are sensitive to alterations in small GTPases such as Rab4, Rab5, Rab8 and Rab11, which regulate endosomal compartments (Callejo et al., 2011; D’Angelo et al., 2015; Gore et al., 2021; Pizette et al., 2021; Stewart et al., 2018). The expression of Disp-1-CS previously suggested a requirement of Disp-1 cleavage in endosomal membrane trafficking (Stewart et al., 2018). More recent evidence in *Drosophila* highlights a role of Disp and Rab11 in Hh delivery from a tubular endosomal compartment named Hherisomes, which modifies Hh signaling potential (Pizette et al., 2021). It is unknown whether this compartment is functionally related to the Rab11-ARE compartment of MDCK cells (Perez Bay et al., 2016). Our microinjection experiments show colocalization of newly synthesized Disp-1 and Shh together with Rab11 in a tubular subapical endosome corresponding to the ARE. Furthermore, we show that Disp-1-CS not only decreases basolateral secretion from the TGN but also accumulates Shh at the ARE, thus decreasing its apical cell surface distribution. Therefore, our results add more evidence to support a role of Disp-1 in the vesicular trafficking of Hhs, revealing a requirement for Shh trafficking along the basolateral-to-apical transcytosis pathway at the level of the Rab11-ARE compartment.

The complex trafficking behavior of Shh in MDCK cells is compatible with the apical and/or basolateral distribution and secretion of Hh seen in different epithelial cells, predicting also a reciprocal basolateral versus apical secretion variations. In MDCK cells, the predominant Shh apical location and secretion at steady state would result from low levels of Shh basolateral secretion. Parietal cells in the stomach distribute and secrete Shh mostly apically, also showing detectable levels at the basolateral side (Zavros et al., 2008). Basolateral secretion of Shh can be inferred from stromal mesenchymal cell responses in intestinal cells (Buller et al., 2012; Shyer et al., 2015) and kidney epithelium (Ding et al., 2012; Edeling et al., 2016; Zhou et al., 2014), without discarding an apical secretion with other functions. Apical and basolateral secretion of Shh sustaining distinct roles and signaling have been described in mature airway epithelia (Mao et al., 2018). Here, immunostaining localizes Shh both at apical and lateral regions and Shh basolateral release stimulates the stromal mesenchymal cells through primary cilia, while its apical secretion induces non-canonical signaling via motile cilia (Mao et al., 2018). In our proposed model the efficiency of Shh endocytic removal for transcytosis and/or secretion from the basolateral domain would determine the relative distribution and secretion from each domain.

Our results might also help to reconcile debated models in *Drosophila* epithelial systems (Guerrero and Kornberg, 2014; Matusek et al., 2020). The relative magnitude of apical and basolateral exocytic pathways operating in the long studied wing disc epithelia is unknown (Guerrero and Kornberg, 2014; Matusek et al., 2020). Extensive studies in the wing disc epithelia proposes a primary addressing of Hh towards the apical plasma membrane, even though it is detected both in apical and basolateral cell surfaces (Callejo et al., 2011; D’Angelo et al., 2015; Gradilla and Guerrero, 2013; Therond, 2012). A long controversy stands on whether secretion for long-range Hh signaling occurs from the apical domain after endocytic recycling (D’Angelo et al., 2015), or from the basolateral cell surface following apical-to-basolateral transcytosis (Callejo et al., 2011). However, previous studies suggested a primary Hh basolateral addressing followed by transcytosis to the apical domain instead, which might explain the enhanced basolateral Hh distribution seen when Disp is silenced in other embryo epithelial cells (Gallet et al., 2003). Basolateral Hh accumulation and secretion has also been found under conditions that silenced the small GTPase Rab8, known to be involved in endocytic and basolateral trafficking (Gore et al., 2021). These controversial views reflect the difficulties of attributing a trafficking pathway based on the interpretation of locations and cell target responses under steady-state conditions. As we show in MDCK cells, a predominant steady-state apical cell surface distribution can result from a primary basolateral sorting from the TGN followed by transcytosis. Our results also demonstrate that transcytosis of Shh still bound to the membrane occurs not only from the basolateral but also from the apical domain. Although we did not measure apical recycling after apical endocytosis, as described in the wing disk epithelia (D’Angelo et al., 2015), this pathway might indeed operate together with transcytosis. Therefore, our results are compatible with the observations in *Drosophila* epithelial systems and perhaps can be used to reinterpret conflicting Hh trafficking models.

Hh proteins have been found secreted in different ways, including multimers (Gallet et al., 2006; Zeng et al., 2001), lipoprotein particles (Panakova et al., 2005), extracellular vesicles (Gradilla et al., 2014; Matusek et al., 2014; Parchure et al., 2015; Tanaka et al., 2005; Vyas et al., 2014) and associated with cytonemes that are long filopodia (Bischoff et al., 2013; Chen et al., 2017; Gonzalez-Mendez et al., 2017). How a primary basolateral sorting followed by transcytosis relates with particular forms of Shh secretion remains a challenge for future studies. Hh proteins form nano-scaled oligomers via electrostatic interactions and this process is required for endocytosis followed by targeting to multivesicular bodies and secretion in exosomes (Parchure et al., 2015; Vyas et al., 2008; Vyas et al., 2014). The role of this Shh oligomerization in polarized sorting can now be evaluated in future experiments using the proposed model in MDCK cells.

An indirect trafficking towards the apical cell surface includes several intermediate stages where regulation might occur impacting upon Shh polarized secretion. This would allow for reciprocal basolateral versus apical secretion variations in distinct epithelial contexts. Conditions that increase basolateral secretion would decrease apical secretion. Variations in Disp-1 expression will first impact the Shh basolateral secretion and then its transcytosis-dependent apical secretion. This, together with the regulation of endocytic trafficking through Rab GTPases, can contribute to determine variations in the basolateral versus apical secretion and the signaling extension of this powerful regulatory molecule in morphogenesis and cancer.

## MATERIALS AND METHODS

### Experimental procedures

#### Key resources table

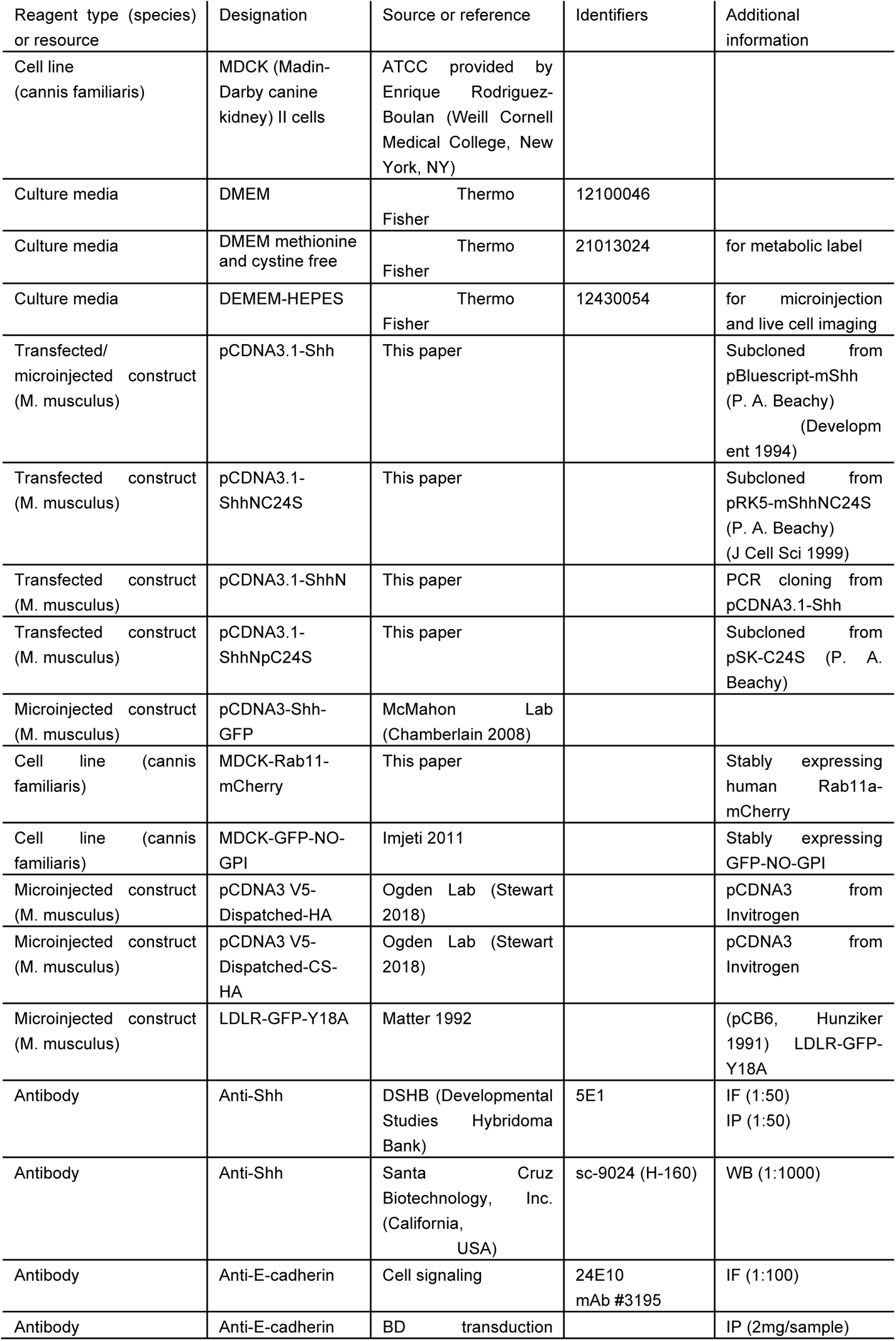
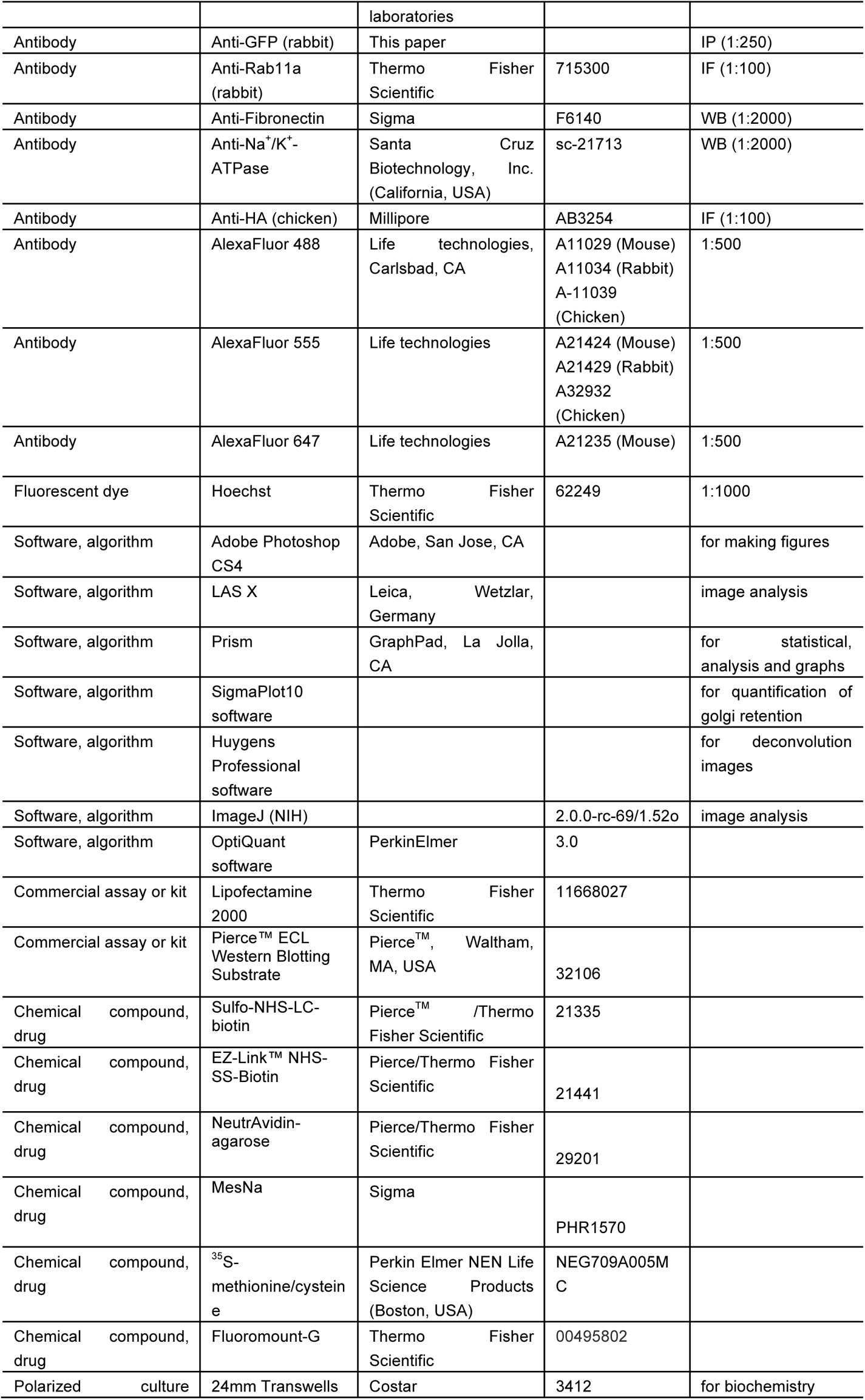

#### Cloning and Subcloning

cDNA coding for Shh (from pBlueScript-Shh), ShhNpC24S (from pSK-C24S) and ShhNC24S (from pRK5-ShhNC24S) (kindly provided by Philip A. Beachy, Stanford University, CA, USA) were subcloned by double digestion with EcoRI and XbaI (for Shh), NotI (for ShhNpC24S) or NotI and KpnI (for ShhNC24S) restriction enzymes and ligated into pcDNA3.1(-) with T4 DNA ligase (Thermo Fisher cat #15224017) according to the manufacturer’s instructions. ShhN dsDNA coding for residues 1-198 of murine Shh was obtained by PCR using Pfu polymerase (Promega cat # M7741) and the following primers: Fw: GTCTCGAGCCAACTCCGATGTGTTCCG and Rev: CGGAATTCTTAGCCGCCGGATTTGGCC that include XhoI and EcoRI sites for subcloning into pcDNA3.1(+). All constructs were confirmed by sequencing.

#### Cell Lines

MDCK cells were grown in DMEM (Cat# 12100046, Thermo Fisher) supplemented with 7.5% fetal bovine serum (FBS, Cat# SH30071.03 HyClone), 50 U/mL antibiotics (Penicillin-Streptomycin mix (Cat# 15070063, Thermo Fisher), and 5 µg/ml Plasmocin (Cat# ant-mpp, InvivoGen), in a 37°C, 5% CO_2_ humidified incubator, routinely analyzed by RT-PCR for mycoplasma contamination. Passages were made before reaching confluence every 2-3 days by two washes with PBS (15 min each) and incubation in trypsin 0.05% for 5-10 min (Trypsin Cat#15400054, Thermo Fisher). MDCK cells stable transfected clones (MDCK-Shh, -ShhN, -ShhNC24S, -ShhNpC24S) were generated by transfection with Lipofectamine 2000 reagent and selection by G418 resistance.

#### Transfection

MDCK (100.000 cells) grown in 24-multiwell plates were transfected with 1µg of plasmid DNA and Lipofectamine 2000 reagent, trypsinized the next day and plated at 1/10 and 1/100 dilution into 100mm dishes. Selection media containing G418 (0.8mg/ml) was added the next day, changed every 5 days until visible colonies were isolated using cloning cylinders and expanded in 6-multiwell plates. Screening was made by immunofluorescence (see below) and immunoblot of lysates and media (See “Protein Analysis”).

#### Cell lysates and protein analysis

To obtain lysates cells were washed twice in PBS containing 1 mM MgCl2, 0.1mM CaCl2 (PBS-CM) and lysed in 500 µl RIPA buffer (50 mM Tris-HCl (pH 8.0), 150 mM NaCl, 0.5% Na-deoxycholate, 0.1% SDS, 1% Triton X-100) supplemented with protease inhibitors (PMSF 2mM, Leupeptin 2µg/ml, Pepstatin 1µg/ml). Recovered lysates were cleared by centrifugation at 14.000rpm at 4°C. Protein concentration was determined on the cleared cell lysates with BCA method and 500 µg total protein was used for immunoprecipitation or NeutrAvidin-precipitation (see below), 10% volume of each lysate was used as loading control.

#### Steady-state Shh Secretion

Analysis of secreted proteins was performed by collection of conditioned media in low serum media (0.3% FBS), precipitation in 10% trichloroacetic acid (TCA) and incubation for 30min on ice. Precipitated proteins were centrifuged at 4°C and 14.000 rpm for 10 min, washed once with 1ml of 100% acetone and re-centrifuged. Pellets were solubilized in 30μl of NaOH 0.1M and 10μl of 3X Laemmli sample buffer. Lysates and/or secreted proteins were resolved in 12%SDS-PAGE, transferred to PVDF membranes and immunoblotted with the indicated antibodies.

#### Polarized monolayers of MDCK cells grown in Transwell inserts

MDCK cells (100,000-400,000 cells) were plated on 12 mm or 24-mm Transwell inserts, 0.4-mm pore size (Costar), and grown with daily media changes until the transepithelial resistance reached 400 ohm/cm^2^ (4–5 days) measured with an EVOM electrometer (World Precision Instruments, Sarasota, FL).

#### Immunofluorescence

Cells were fixed in freshly prepared 4% PFA in PBS-CM for 15 min at room temperature or overnight at 4°C. To analyze subcellular distribution of interest proteins, cells were permeabilized with 0.2% saponin in PBS-CM. All subsequent steps contained 0.2% saponin. Primary and secondary antibodies were incubated for 30 min at 37°C or overnight at 4°C. Coverslips were mounted in Fluoromount.

#### Protein immunoprecipitation

The key resources table indicates the amount of antibodies (GFP, E-cadherin, 5E1) used per sample. Antibodies were pre-incubated with 30µl of protein-A sepharose for 2h at 4°C, then beads were washed 3 times to remove excess antibodies. Total cell lysate (approx. 500 µg) or collected medium was then added to beads and incubated for 2h at 4°C, washed 3 times in RIPA buffer and eluted in 30µl of 2X Laemmli Buffer.

#### Domain-Selective Surface Biotinylation

Steady-state apical and basolateral distribution of Shh and endogenous proteins were analyzed by domain selective biotinylation of MDCK cells grown in Transwell inserts, as described (Bravo-Zehnder et al., 2000; Oyanadel et al., 2018). Cells were rinsed twice with ice-cold PBS-CM followed by two successive 20 min incubations at 4°C with 0.5 mg/ml Sulfo-NHS-LC-Biotin in PBS-CM, added to the apical (0.6 ml) or basolateral (1 ml) sides. Cells were then incubated with 50mM NH_4_Cl in PBS-CM for 10 min, rinsed twice with PBS-CM and lysed in RIPA buffer, supplemented with proteases inhibitors for 30 min at 4°C. NeutrAvidin-Sepharose was used to retrieve biotinylated proteins by centrifugation (18,000 x g for 10 min), which then were analyzed by immunoblot against Shh and endogenous proteins, as described (Bravo-Zehnder et al., 2000; Oyanadel et al., 2018; Sargiacomo et al., 1989).

#### Pulse-chase domain-selective targeting of newly synthesized proteins

Biotinylation targeting assay was used to assess biosynthetic delivery of proteins to apical and basolateral PM domains. Stable transfected cells were pulse-labeled in methionine-free and cysteine-free DMEM supplemented with 400 µCi/ml [^35^S]-methionine/cysteine for 30 min in 120µl of medium applied to the basolateral side of the inverted filter, as described (Marzolo et al., 1997). Chase was made for different times in culture media containing 300 µM cycloheximide (CHX) and 3- to 6-fold excess cold methionine (0.6mM) and cysteine and (1.2mM). Cells were then subjected to domain-selective surface biotinylation, chilled in ice-cold PBS-CM and lysed with RIPA buffer supplemented with protease inhibitors at 4°C for 30 min. Lysates were cleared by centrifugation (18,000 x g for 10 min) before immunoprecipitation with protein-A-Sepharose preincubated antibodies (5E1-anti-Shh or E-cadherin). The mixture of biotinylated (surface) plus non-biotinylated (intracellular) immunoprecipitated proteins was incubated at 95° for 5 min in 30 µl of 10% SDS RIPA to dissociate immune complexes. The eluted proteins were diluted to 1 ml with RIPA and subjected to NeutrAvidin-Sepharose precipitation to isolate surface-located Shh. In addition, secreted Shh was immunoprecipitated from the apical and basolateral media with protein-A-Sepharose bound 5E1, as described for other proteins (Gonzalez et al., 1993; Gonzalez et al., 1987; Marzolo et al., 1997). The immune complexes of surface-located and secreted Shh were resolved by SDS-PAGE. [^35^S]-methionine/cysteine labeled proteins were visualized in fluorograms made with Amplify^TM^ fluorography reagent (Amersham), developed in scanner Cyclone Plus (Perkin Elmer) with OptiQuant software (3.0, Perkin Elmer).

#### Transcytosis assays

Trafficking of Shh from one cell surface domain to the opposite (transcytosis) was analyzed by domain-specific biotinylation and cell surface-tagging immunofluorescence assays.

*1. Biotinylation transcytosis assay* was performed with reducible biotin reagent Sulfo-NHS-SS-biotin as described (Burgos et al., 2004). In brief, six Transwell filter cups with polarized live MDCK monolayer cells were subjected to basolateral-specific biotinylation with Sulfo-NHS-SS-biotin in *PBS-CM.* One filtercup was used for total basolateral biotinylated Shh (“non-reduced, time 0”), and other two filtercups were subjected to either apical or basolateral reduction with 22 mM MesNa reagent (0 min) made in buffer 50 mM Tris-HCl, pH 8.6, 100 mM NaCl. The remaining three filtercups were incubated for 120 min in 37°C in growth media to allow trafficking and then MesNa was added to the apical or basolateral side. Monolayers were alkylated with 20mM iodoacetamide in PBS-CM containing 1%BSA, washed twice in PBS-CM, lysed in Ripa Buffer supplemented with protease inhibitors and subjected to NeutrAvidin-Sepharose precipitation and immunoblot. To analyze apical secretion of Shh present in the basolateral membrane, cells were biotinylated from the basolateral side and then incubated in DMEM, 0.5 % fetal bovine serum at 37°C for 2 hours. Biotinylated Shh proteins were retrieved from apical and basolateral media with NeutrAvidin-Sepharose and immunoblotted.
*2. Cell surface-tagging immunofluorescence transcytosis assay.* Live MDCK cells grown in Transwell inserts were apically or basolaterally incubated with 5E1 antibody diluted in DMEM (supplemented with 10mM HEPES) for 30 min at 4°C. After 3 washes in PBS-CM, cells were shifted to 37°C in growth media to allow trafficking for different times, fixed at different times with PFA-PBS-CM and permeabilized. Shh bound to the primary antibody was detected by incubation of fluorophore-coupled secondary antibodies for 30 min at 37°C.
*3. Secretion of basolaterally-sorted ShhNpC24S.* MDCK cells grown in Transwell inserts were rinsed twice with ice-cold PBS-CM followed by two successive 20 min incubations at 4°C with 0.5 mg/ml Sulfo-NHS-LC-Biotin in PBS-CM, added to the basolateral sides. Cells were then incubated with 50mM NH_4_Cl in PBS-CM for 5 min, rinsed twice with PBS-CM and shifted to 37°C in growth media for 2h. Then apical and basolateral media was collected and precipitated with Neutravidin (basolaterally-biotinylated secreted proteins). Samples were analyzed by western blot.

#### Microinjection

MDCK cells were plated either in 12mm diameter glass coverslips (5000 cells) the absence of Plasmocin. After achieving full confluence (4-5 cells) microinjections were performed in the cell nucleus (plasmids) using back-loaded glass capillaries, an Eppendorf NI2 micromanipulator and an Eppendorf Transjector 5246 system. The expression plasmids DNAs were used at a concentration of 20 to 100ng/µl in buffer HEPES 10 mM pH7.4, KCl 140 mM. During the microinjection procedure cells were maintained in DMEM-HEPES.

#### Live cell imaging

The microinjection approach to express and accumulate exogenous proteins at the TGN and then assess subsequent trafficking was performed as described (Cancino et al., 2007; Kreitzer et al., 2000). Briefly, microinjected cells were incubated for 45 min at 37°C in DMEM-HEPES plus 7.5% FSB to allow protein expression without ER leakage, plasmid concentration was adjusted (20 to 100ng/µl) to obtain enough protein expression to be detected by fluorescence microscopy and then the exocytic trafficking was arrested at the TGN by incubating the cells at 20°C for 2 h in DMEM-HEPES with CHX. After the 20°C block the cells were mounted on a Leica SP8 microscope in DMEM-HEPES plus CXM and incubated at 37°C.

#### Images acquisition and processing

Images were collected at 1024 × 1024 resolution in Z stacks of 300-nm steps using a Leica SP8 confocal microscope and a 63× oil immersion 1.4 ma lens, and processed with ImageJ free software, NIH. High Resolution images were acquired at Nyquist rate parameters (Nyquist Calculator, SVI, NL) and the resulting oversampled images were deconvoluted with Huygens Essential Software (SVI, NL), which was also used for 3D surface rendering processing.

#### Statistical Analysis

Significance was determined with Prism software using Student’s t-test. For all statistical analyses **p<0.05 and ****p<0.0001. Error bars indicate standard error of mean (SEM).

## ACKNOWLEDGEMENTS

We are grateful to Dr. Stacey Ogden (St. Jude Children’s Research Hospital, USA) for kindly providing Dispatched plasmids constructs, Dr. Andrew P. McMahon (University of Southern California, USA) and Dr. Philip A. Beachy (Stanford University School of Medicine, USA) for kindly providing Shh plasmids constructs.

## ADDITIONAL INFORMATION

## Funding

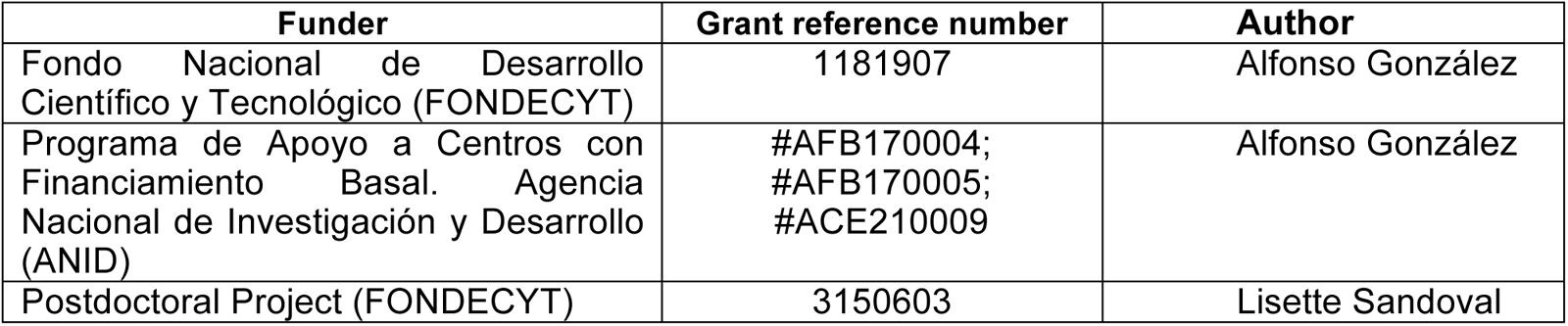

## AUTHOR CONTRIBUTIONS

Lisette Sandoval, Conceptualization, Data curation, Formal analysis, Validation, Investigation, Visualization, Methodology, Writing - original draft, Writing - review and editing; Mariana Labarca, Conceptualization, Formal analysis, Validation, Investigation, Visualization, Methodology, Writing - review and editing; Claudio Retamal, Data curation, Formal analysis, Methodology; Juan Larraín, Conceptualization, Resources; Alfonso González, Conceptualization, Supervision, Funding acquisition, Writing

